# Maximum entropy models reveal the correlation structure in cortical neural activity during wakefulness and sleep

**DOI:** 10.1101/243857

**Authors:** Trang-Anh Nghiem, Bartosz Telenczuk, Olivier Marre, Alain Destexhe, Ulisse Ferrari

## Abstract

Maximum Entropy models can be inferred from large data-sets to uncover how local interactions generate collective dynamics. Here, we employ such models to investigate the characteristics of neurons recorded by multielectrode arrays in the cortex of human and monkey throughout states of wakefulness and sleep. Taking advantage of the separation of excitatory and inhibitory types, we construct a model including this distinction. By comparing the performances of Maximum Entropy models at predicting neural activity in wakefulness and deep sleep, we identify the dominant interactions between neurons in each brain state. We find that during wakefulness, dominant functional interactions are pairwise while during sleep, interactions are population-wide. In particular, inhibitory neurons are shown to be strongly tuned to the inhibitory population. This shows that Maximum Entropy models can be useful to analyze data-sets with excitatory and inhibitory neurons, and can reveal the role of inhibitory neurons in organizing coherent dynamics in cerebral cortex.

## I. INTRODUCTION

To analyze a complex system, one is interested in finding a model able to explain the most about empirical data, with the fewest forms of interactions involved. Such a model should reproduce the statistics observed in the data, while making the least possible number of assumptions on the structure and parameters of the system. In other terms, one needs the simplest, most generic model that generates statistics matching the empirical values - this implies *maximisation of entropy* (MaxEnt) in the system, with constraints imposed by the empirical statistics [1].

In a seminal paper [2], a framework equivalent to the Ising model in statistical physics was used to analyze the collective behavior of neurons. This approach was based on the assumption that pairwise interactions between neurons can account for the collective activity of the neural population. Indeed, it was shown for experimental data, from the retina and cerebral cortex, that this approach can predict higher order statistics, including the probability distribution of the whole population’s spiking activity. Even though the empirical pairwise correlations were very weak, the model performed significantly better than a model reproducing only the firing rates without considering correlations. The Ising model was subsequently demonstrated to efficiently reproduce the data better than models with smaller entropy [3], as well as to analyse neural recordings in a variety of brain regions in different animals, ranging from the salamander retina [2, 4] to the cerebral These authors contributed equally. cortex of mice [5], rats [6], and cats [7].

A complementary approach was recently introduced [8], aiming at reproducing the correlation between single neuron activity and whole-population dynamics in the mouse and monkey visual cortex. Later work [9] generalised this approach to model the neurons’ full profile of dependency with the population activity, and applied the model to the salamander retina.

Recent advances in experimental methods have allowed the recording of the spiking activity of up to a hundred neurons throughout hours of wakefulness and sleep, for instance using multi-electrode arrays, also known as Utah arrays. Inspection of neurons’ spike waveforms and their cross-correlograms with other neurons made the discrimination of excitatory (E) and inhibitory (I) neuron types possible [10, 11]. Such data-sets therefore provide a further step in the probing of the system, due to the unprecedented availability of the simultaneously recorded dynamics of E and I neurons.

In the present paper, we apply Maximum Entropy models to analyze human and monkey Utah array recordings, which is done here for the first time. We also investigate in which way such models may describe the two recorded (E, I) populations, and what supplementary insight they provide on the interplay between excitation and inhibition.

## II. RESULTS

We study 96-electrode recordings (Utah array) of spiking activity in the temporal cortex of a human patient and in the premotor cortex of a macaque monkey (see Appendix A), in wakefulness and slow-wave sleep (SWS), as shown in Fig. 1. Spike times of single neuronal neurons were discriminated and binned into time-bins of 50 ms (human data) and 25 ms (monkey data) to produce the population’s spiking patterns (see Appendix A). From these patterns, we computed the empirical covariances between neurons then used for fitting models.

**FIG. 1:**
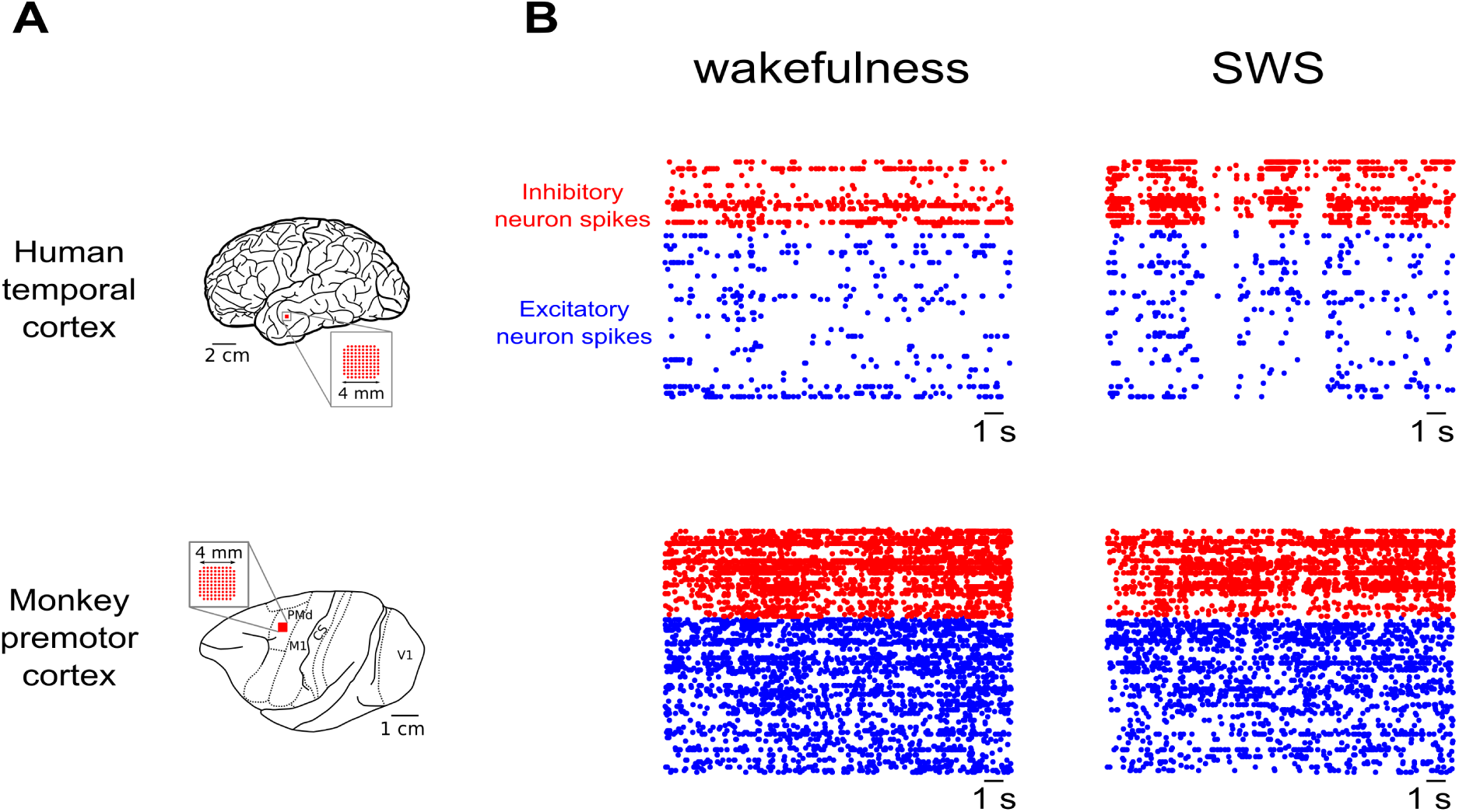
Multi-electrode (Utah) array recordings. **A)** Utah array position in human temporal cortex (top) and monkey prefrontal cortex (bottom). Figure adapted from [12]. **B)** Raster plots of spikes recorded for human (top) and monkey (bottom) in wakefulness (left) and SWS (right). Excitatory (E) neurons are represented in blue, while inhibitory (I) neurons are in red.

### Pairwise Ising model

Pairwise correlations between I neurons have been found to exhibit invariance with distance [13], even across brain regions [14]. Here, we study what this intriguing observation implies for functional interactions between neurons, and the information conveyed by pairwise correlations on such interactions. Therefore, we investigate whether pairwise covariances are sufficient to capture the main features of neural activity, for E and I neurons during wakefulness and SWS.

To test this, we use a MaxEnt model that reproduces only and exactly the single neurons’ spiking probability, and the pairwise covariances observed in the data.

As it has been shown [2, 15], this model takes the form of a disordered Ising model (see Fig. 2A):

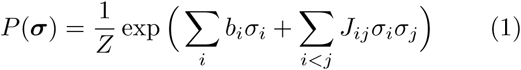

where σ_*i*_ denotes activity of neuron i given time bin (1: spike, 0: silence), b_*i*_ the bias of neuron *i*, controlling its firing rate, and *J*_*ij*_ the (symmetric) coupling between neurons *i* and *j*, controlling the pairwise covariance between the neurons.

**FIG. 2:**
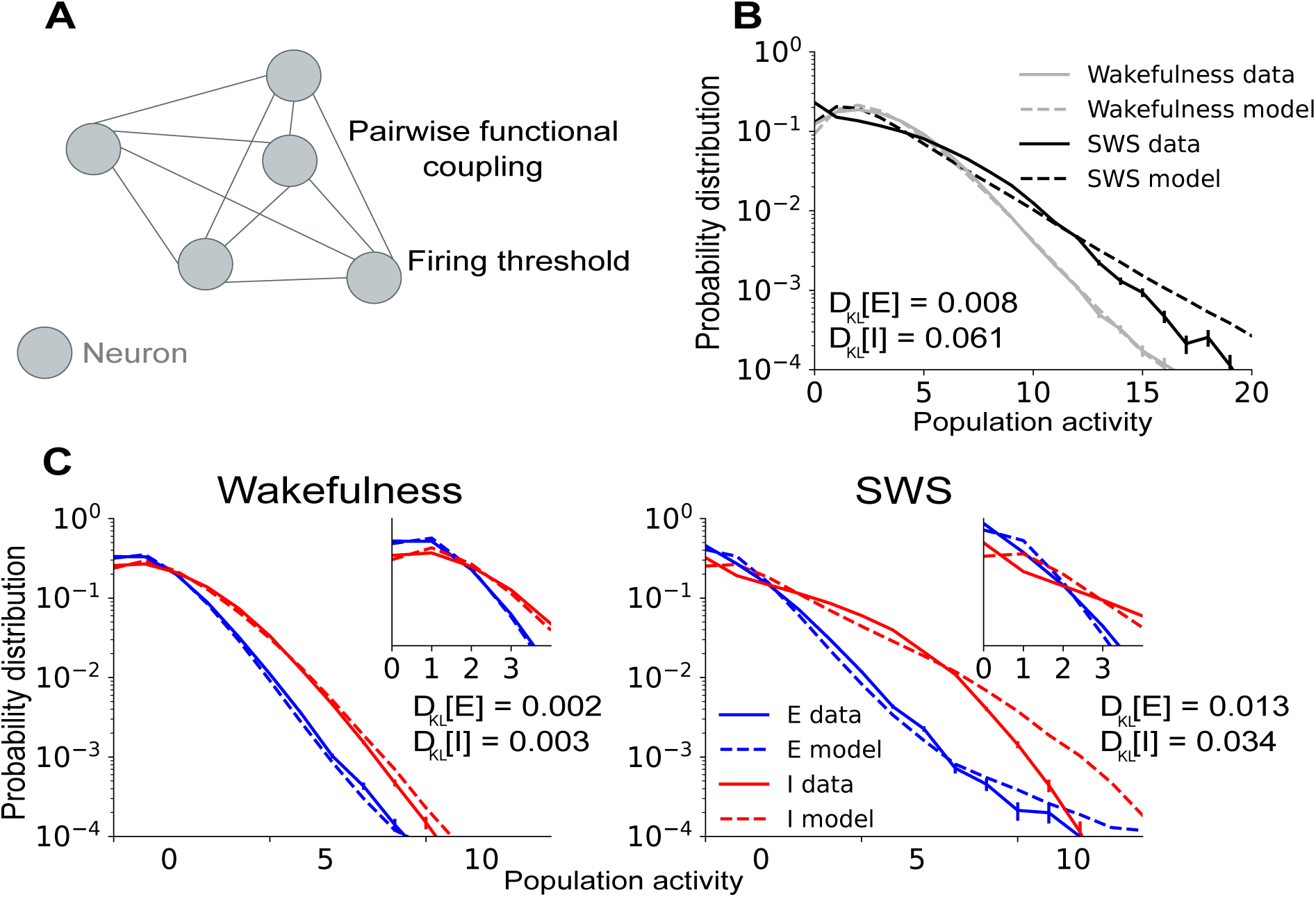
Pairwise Ising model fails to predict SWS synchronous activity, especially for inhibitory neurons. **A**) Model schematic diagram. Model parameters are each neuron’s bias toward firing, and symmetric functional couplings between each pair of neurons. **B**) Empirical and predicted probability distributions of the population activity 
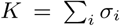
 for the population of excitatory (E) neurons, and that of inhibitory (I) neurons. The Ising model more successfully captures the population statistics during wakefulness than SWS, especially for medium and large *K* values. **C**) Empirical and predicted population activities for E and I neurons. The model particularly fails at reproducing the statistics of I population activity. These results are consistent with the presence of transients of high activity and strong synchrony between I neurons during SWS.

We use the algorithm introduced by [16] to infer the model’s parameters *b*_*i*_ and *J*_*ij*_ on data from wakeful-ness and SWS separately, for each Utah array recording. Then we test how well the model describes neural activity in these states. In particular, synchronous events involving many neurons may not be well accounted for by the pairwise nature of the Ising model interactions. To test this, we quantify the empirical probability of having *K* neurons active in the same time window [4]: 
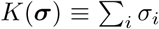

Fig. 2B compares the empirical probability distributions with model predictions. The Ising model is able to account for the empirical statistics during wakefulness, while it partially fails to capture the statistics during SWS. This is confirmed by the measures of the Kullback-Leibler divergence, 
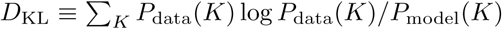
 between empirical and model-predicted distributions (Fig. 2B). This difference can be ascribed to the presence of high activity transients, known to modulate neurons activity during SWS [17] and responsible for the larger covariances, as seen in [10]. In order to investigate the Ising model’s failure during SWS, in Fig. 2C we compare the predictions for *P* (*K*), separating excitatory and inhibitory neuron populations. For periods of wakefulness, the model is able to reproduce both neuron types’ behaviors. However, during SWS periods, the model largely fails at predicting the empirical statistics, in particular for the I population. This is confirmed by estimates of the Kullback-Leibler divergences (see Fig. 2). Fig. S1 shows similar results for the analysis on monkey recording.

These results highlight the relevance of the pairwise Ising model to reproduce *P*(*K*) for all neurons, E and I, during wakefulness. Neural dynamics during wakefulness can therefore be described as predominantly driven by pairwise interactions. However, during SWS the model fails to reproduce *P*(*K*) for both populations. Therefore pairwise couplings alone are not sufficient and higher-order, perhaps even population-wide interactions may be needed to accurately depict neural activity during SWS. This is consistent with the observation that during SWS, neural firing is synchronous even across long distances, most notably for pairs of I neurons [14].

So far, our findings from inferring a pairwise Ising model on our datasets have highlighted that pairwise interactions were sufficient to depict neural activity during wakefulness, but higher-order, population-wide interactions may appear during SWS.

### Single-population model

In order to further characterize the neuronal activity during SWS, we consider the interaction between each neuron and the whole population: indeed, such approaches have proven successful in describing cortical neural activity [8]. We investigate whether neuron-to-population interactions exist in our data-set by studying the neurons’ tuning curves to the population. Neuron-to-population tuning curves (see Appendix C) indicate how much a neuron’s activity is modulated by the total activity of the rest of the network [9]. In Fig. 3A we present tuning curves for ten example E or I neurons during both wakefulness and SWS. These examples provide strong evidence for neuron-to-population tuning. In order to quantify population tuning, we estimate how much a neuron, either E or I, is *sensitive* to the activity of the rest of the population, i.e. how much its activity fluctuates depending on the population activity (see Methods). As can be observed in Fig. 3B, and consistently with our previous results, we find that neurons are sensitive to the population especially during SWS. Similar results are valid for the monkey recording as well (Fig. S2A). Since we have established neuron-to-population interactions take place during SWS, we wish to determine to what extent they are sufficient in capturing the characteristics of neural activity during sleep.

**FIG. 3:**
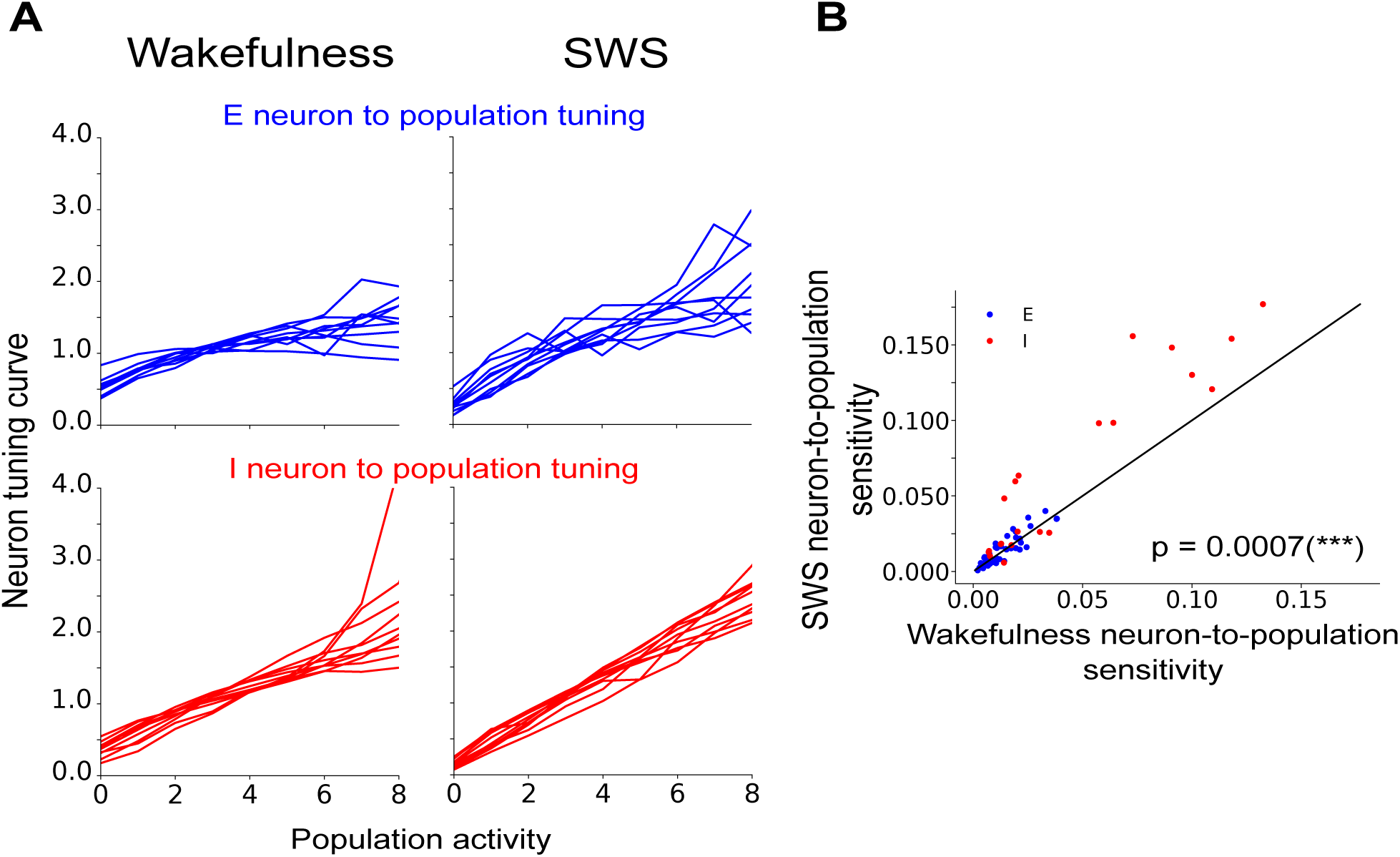
Neural firing is tuned to the neural population’s activity, particularly during SWS. **A**) Tuning curves of ten example neurons (see text and Appendix C) showing that neurons are tuned to the population activity. **B**) Scatter-plot of the neuron sensitivity to the population activity (see Appendix C). Neurons are very consistently more sensitive during SWS (*p*-value < 0:001, Wilcoxon sing-ranked test).

To this purpose, we use a null model [9] for the dependencies between neuron firing, 
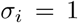
, and population activity, *k*: 
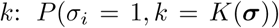
, where 
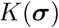
 denotes the number of neurons spiking in any time bin. In this model (Fig. 4A), the probability of neuron ring is described by the strength of its coupling to the population: 

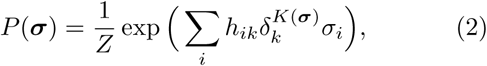

**FIG. 4:**
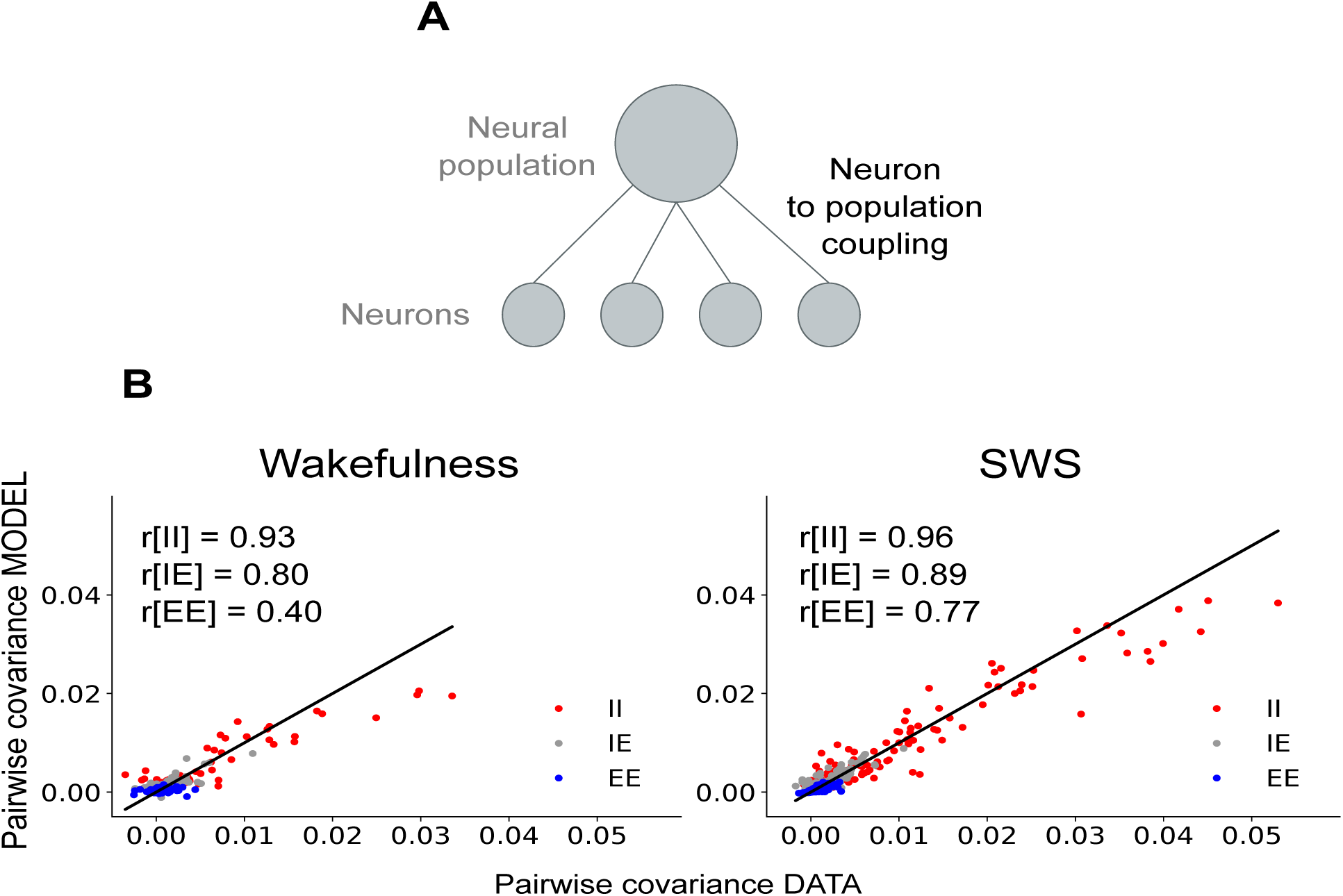
Single-population model shows better performance during SWS than wakefulness. **A**) Model schematic diagram. **B**) Pairwise covariances, empirical against predicted, for wakefulness (left) and SWS (right) states. Consistently with Fig. 3B, the success for SWS, most noticeably for I-I pairs, suggests these neurons are most responsive to whole-population activity.

where *h*_*ik*_ is the coupling between neuron *i* and the whole population when *k* neurons are active. *δ*_*k*_^*k*^ is the Kronecker delta, taking value one when the number *K* of active neurons is equal to a given value *k* and zero otherwise. For example, a \chorister" neu ron, that fires most often when many others are firing, would have *h*_*ik*_ increasing with *k*. Conversely, a “soloist” neuron, that fires more frequently when others are silent, would have *h*_*ik*_ decreasing with *k* [8]. *Z* is the normalisation constant, that can be computed by summing over all possible activity configurations

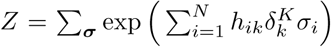
. Importantly, *Z* and its derivative allow us to determine the statistics of the model, such as the mean firing rate and the pairwise covariances. As an analytical expression exists for *Z*, the statistics may be derived analytically from the values of the couplings, making this model solvable (see Appendix).

To evaluate to what extent the model describes the data well and hence captures empirical statistics it was not designed to reproduce, we study the predicted pairwise correlations as compared to the empirical ones.

In Fig. 4B, we compare the empirical pairwise covariances to their model predictions. Pearson correlations (covariance between the two empirical and predicted variables, normalized the product of their standard deviations) confirm that the population statistics are better reproduced by the model during SWS than during wakefulness (Fig. 4). For monkey recording, the effect is even larger since the model entirely fails to account for wakefulness pairwise statistics (Fig. S2B). While the effect may be amplified by the fact that the Pearson correlations are larger during SWS, this is the opposite of what was observed for the pairwise Ising model: a model reproducing only empirical neuron-to-population interactions seems adequate at depicting neural dynamics during SWS but not during wakefulness.

In particular, the model best reproduces the empirical statistics during SWS for I neuron pairs. By contrast, E-E pairwise covariances are the most poorly reproduced during wakefulness. This result implies that during SWS, I activity, and to a lesser extent E activity, is dominated by population-wide interactions rather than local pairwise mechanisms, such that a MaxEnt’population model’ is mostly sufficient at capturing the key dynamics. Nevertheless, this model still underestimates the higher I-I pairwise covariances.

### C. Two-population model

Since I neurons are strongly synchronised even across long distances [10, 11], we hypothesise that they could be tuned to the I population only, rather than the whole population. We therefore ask if I neurons are tuned to the I population only. Indeed, as shown in Fig. 5A, examination of the tuning curves of each neuron to the E and the I populations separately revealed homogeneous and strong tuning of I neurons to the I population, compared to tuning of I neurons to the E population or to the whole population (Fig. 3). In order to quantify this effect, we estimated the neuron sensitivity to both populations separately (see Appendix C). The comparison in Fig. 5B suggests I neurons are significantly more sensitive to the activity of I population than the E population. The effect is even larger for monkey recordings (Fig. S3A).

**FIG. 5:**
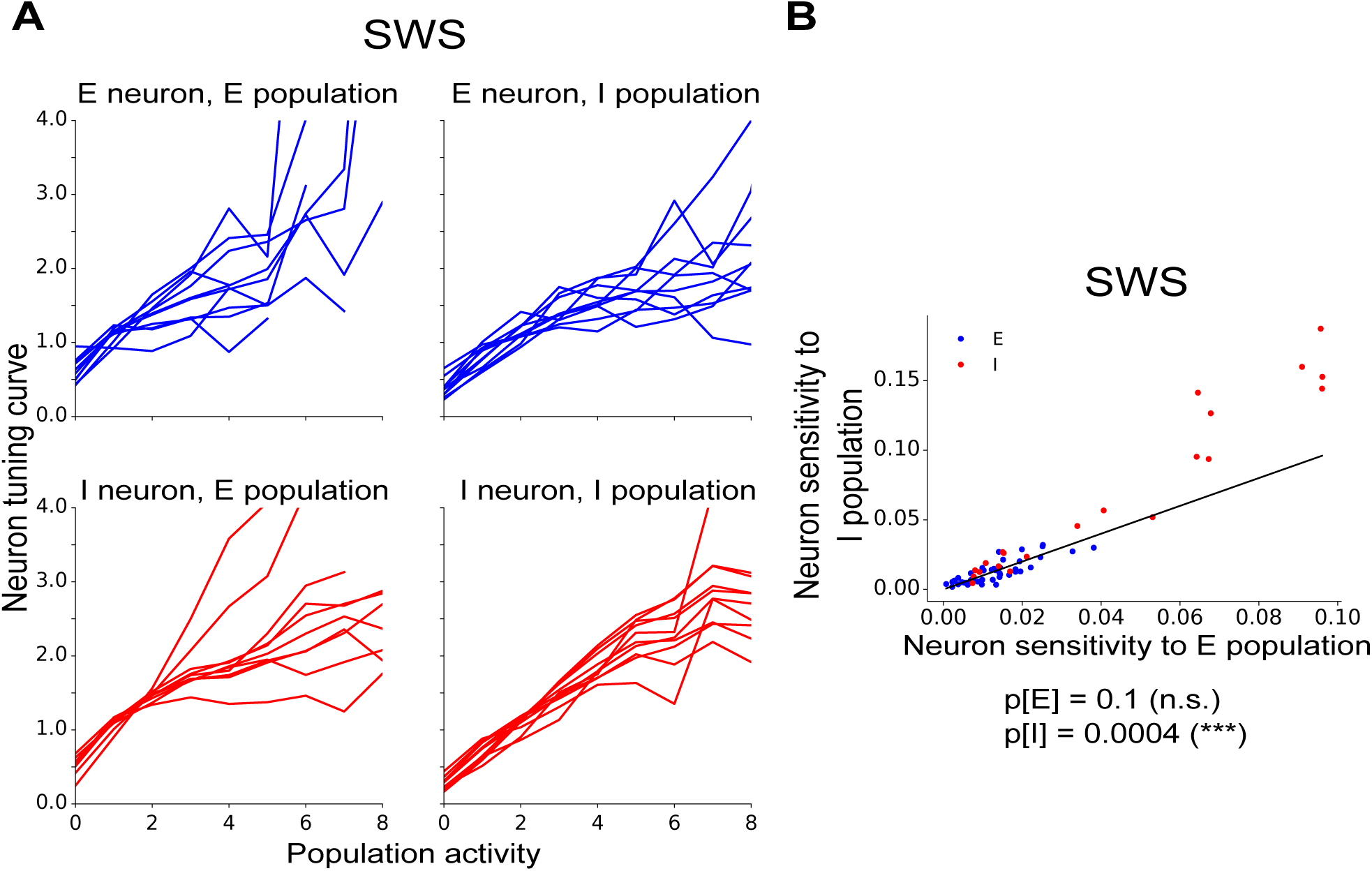
I neurons are more specifically tuned to the I population during SWS. **A**) Example tuning curves from ten neurons of each type to each type of population during SWS, and similarly for the I population. **B**) Scatter-plot of neuron sensitivity to E versus I population, during SWS. I neuron are more tuned to I population than the E population (*p*-value <lt; 10^−3^, Wilcoxon sign-ranked test). E neurons, instead, are weakly sensitive to both populations.

To study tuning to the two populations separately, we now refine the previous model to take into account the couplings between each neuron and the E population and each neuron and the I population, separately. Because of the results of Fig. 5B, we expect this model to perform better at reproducing the main features of the data during SWS. We want the model to only and exactly reproduce the empirical 
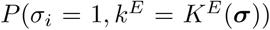

and 
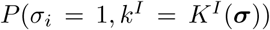
 for all neurons *i* and all values empirically taken by *K*^*E*^ and *K*^*I*^ .

The probability of obtaining any firing pattern σ is given by (see Fig. 6A)

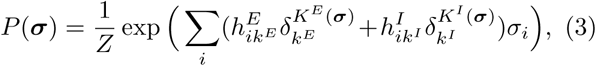

**FIG. 6:**
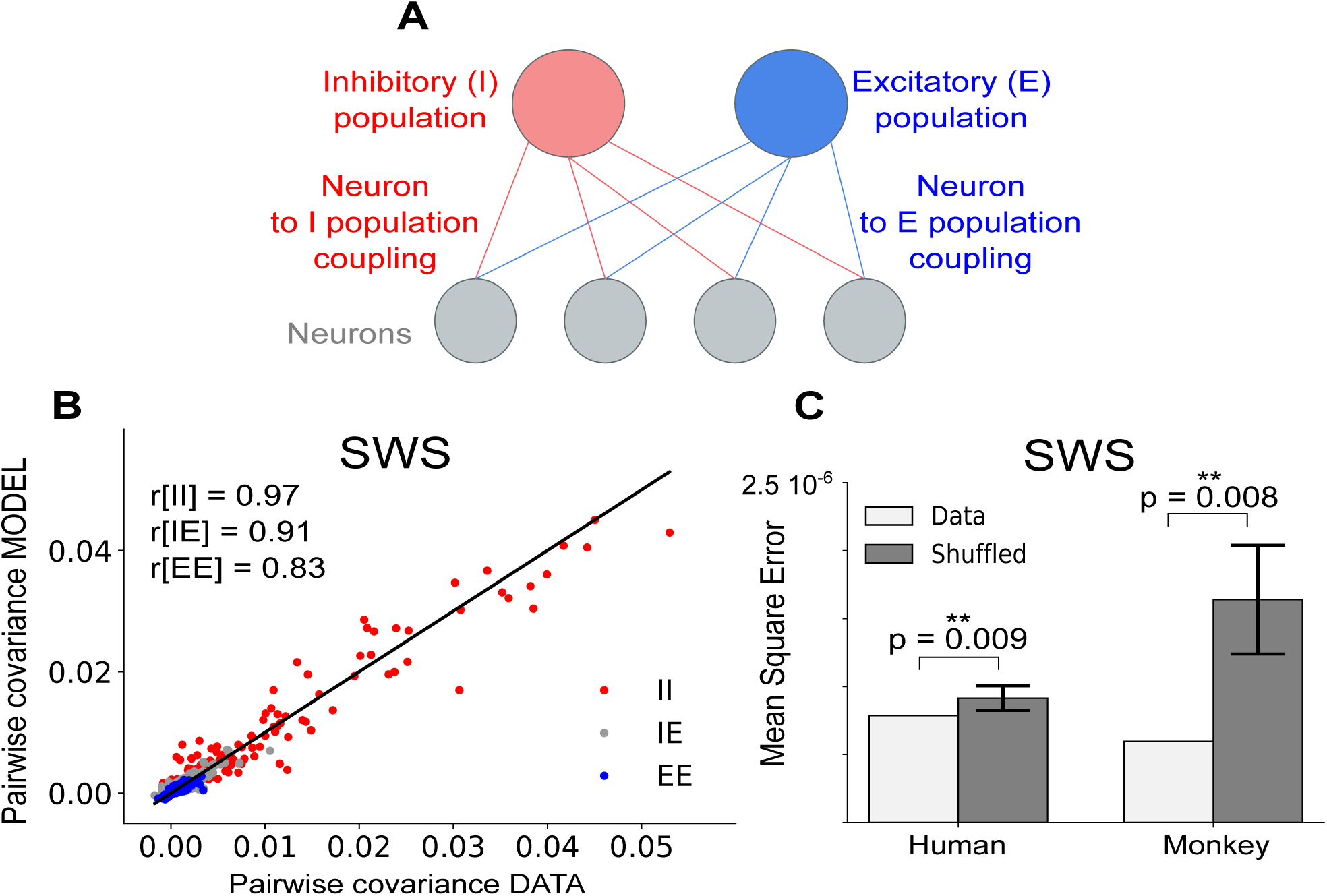
Two-population model shows significant improvement in prediction for all types of neurons. **A**) Schematic diagram of the two population model. Parameters 
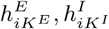
are the couplings between each neuron *i* and the E population activity 
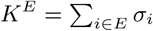
and the I population activity 
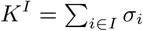
**B**) Pairwise covariances, empirical against predicted, for the two-population model, during SWS. Improvement compared to the whole-population model is confirmed by the Pearson correlations. **C**) deterioration of prediction by shuffling neuron types for the human and monkey data-sets. This e ect demonstrates that knowledge of neuron types signi cantly contributes to improving model prediction. This is con rmed by the Mann-Whitney *U* test *p*-values.

where *K*^E^(*σ*) is the number of E neurons spiking and K^I^ the number of I neurons spiking in any time bin, and 
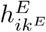
 the coupling between neuron *i* and the whole E population when *k* neurons are active, resp. 
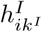
 to the I population. *Z* the normalisation, 
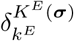
 and

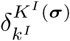
 are Kronecker deltas as before. It can be shown (see Appendix D), following an analogous reasoning to that employed in [9], that this model is also analytically solvable in that the normalisation function *Z* may be derived analytically. Using the expression for *Z*, as described in the Appendix, allows us to analytically predict the model statistics for any given set of couplings. As for the previous models, we want to assess whether this model is sufficient to describe the data, that is if it can accurately predict a data statistic it was not specifically designed to reproduce. To this purpose we test pairwise covariances. We also aim to evaluate how prediction performance compares with the single-population model on the whole population (Fig. 4) described previously.

For both human (Fig. 6B) and monkey (Fig. S3B) recordings, during SWS the two-population model provides better predictions for pairwise covariances than the single-population model. Furthermore large I-I covariance are no longer systematically under-estimated. To verify the improvement in model performance was not solely due to this model possessing more parameters, we repeat the inference on the same data with the neuron types (E or I) shuffled, and find that the prediction deteriorates significantly, as highlighted in Fig. 6C. This demonstrates that taking into consideration each neuron’s couplings with the E population and the I population separately is more relevant than taking into account its couplings with any sub-populations of the same size.

Remarkably, with the two-population model, E-I correlations are also reproduced with increased accuracy as compared to the single-population model. This improvement suggests that the two-population model successfully captures some of the cross-type interactions between the E and I populations, a non-trivial result since the two populations are not directly coupled to one another by design of the model.

## III. DISCUSSION

In this paper, we tested MaxEnt models on human and monkey multi-electrode array recordings where E and I populations were discriminated, during the states of wakefulness and SWS. In order to investigate the properties of the neuronal dynamics, models were designed to reproduce one empirical feature at a time, and tested against remaining statistics. The pairwise Ising model’s performance highlighted pairwise interactions as dominant in cortical activity during wakefulness, but insu cient to describe neural activity during SWS.

We identify I neurons as responsible for breaking pairwise su ciency during SWS, suggesting instead that I neurons’ interactions are long-distance and population-wide, which explains recent empirical observations [10, 14].

We found that models based on neuron-to-population interactions, as introduced by [8], are only relevant to SWS, failing to replicate the empirical pairwise correlations in the monkey premotor cortex (Fig. S2). Even for SWS, I neurons’ strong pairwise correlations were consistently underestimated.

Eventually, the two-population model provides a good trade-off for modelling neural interactions in SWS, and in particular the strongly correlated behaviour of I neurons. Discrimination between E and I neuron types greatly improves the capacity of a model to capture empirical neural dynamics.

### Pairwise sufficiency in Ising MaxEnt models

Pairwise Ising models (Fig. 2A) had previously been shown to accurately predict statistical features of neural interaction in many of data-sets [2, 5, 6, 18]. The surprisingly good performance of these models has raised hypotheses on the existence of some unknown mechanisms beyond their success [19]. In order to understand the so-called ‘pairwise sufficiency’, a number of theoretical investigations [20-23] and an empirical benchmark [3] have been conducted. Model limitations have also been subject to some characterization. For instance, the breakdown of model performance for very large system sizes has been evidenced on experimental data [4] and studied theoretically [24]. Ising model performance has also been shown to be sensitive to time bin size, and to its relation to characteristic time scales of the studied system [25]. Here, we observed that for the same neural system, activity can be well-reproduced in one brain state (wakefulness) and not the other (SWS) (see Fig. 2B). This result reinforces the idea that pairwise su ciency depends on the system’s actual statistical properties, and it is not a more general consequence of the MaxEnt principle.

#### Neurons-to-population couplings

Although our study is the first to propose couplings between neurons and single-type population, an alternative approach has been previously used to highlight the neurons’ tuning by the population activity [8]. In that work, neurons were classified as ‘soloist’ or ‘chorister’, depending on whether they spiked more frequently when the rest of the population was silent or active, respectively. In this work, we have refined this picture by pointing out tuning by single-type population. Specifically, we have shown that I neurons are more sensitive to the I population activity than to the E one (Fig. 5B).

#### Neuron-to-neuron and neuron-to-population couplings as the competition between internal network dynamics and common external inputs.

We note that mechanisms underlying neuronal interactions may occur at multiple scales, as well as involve the interplay between scales. Indeed, population-wide interactions winning over pairwise interactions may be due to a different network connectivity, with reinforced structural couplings between I neurons over long distances. However, this could also be explained by I neurons presenting very similar responses to received inputs from other brain areas (as highlighted by similar tuning curves). Finally, larger common inputs to the I population, may also be a plausible mechanism driving the observed activity.

Biophysical models of spiking neurons could allow for the independent exploration of each of these mechanisms and their effects on the resulting neuron-to-neuron and neuron-to-population interactions. Conversely, MaxEnt models could be used to constrain biophysical models in providing a quantitative way of comparing their statistical features to empirical neural statistics.

## Acknowledgments

We thank C. Capone, M. Chalk, C. Gardella, J.S. Goldman, A. Peyrache, and G. Tkacik for useful discussion. Research funded by European Community (Human Brain Project, H2020-720270), ANR TRAJECTORY, ANR OPTIMA, French State program Investissements d’Avenir managed by the Agence Nationale de la Recherche [LIFESENSES: ANR-10-LABX-65], NIH grant U01NS09050 and a AVIESAN-UNADEV grant.

## Appendix A: DATA-SET

We work with an intra-cranial multi-electrode array recording of 92 neurons in the temporal cortex of an epileptic patient, the same data-set used by [10] and [11]. The record of interest spans across approximately 12 hours, including periods of wakefulness as well as several stages of sleep. Recordings were performed in layer II/III of the middle temporal gyrus, in an epileptic patient (even though far from the epileptic focus and therefore not recording epileptic activity outside of generalised seizures). Data acquisition in that region was enabled by implanting a multi-electrode array, of dimensions 1 mm in thickness and 4x4 mm in area, with 96 micro-electrodes separated by 400*μm* spacings. The array was originally implanted for medical purposes. A 30-kHz sampling frequency was employed for recording. Switches in brain state (wakefulness, SWS, REM, seizure, …) throughout the recording were noted from the patient’s behavioural and physiological parameters. Using spike sorting methods on the obtained data, 92 neurons have been identified. Analysis of the spike waveforms for each of these neurons allowed their classification as putative excitatory (E) and inhibitory (I) neurons. Using the spike times of each neuron, cross-correlograms for all pairs of neurons were also computed to determine whether each neuron’s spikes had an excitatory (positive correlation) or an inhibitory (negative correlation) effect on other neurons through putative monosynaptic connections. It should be noted that neurons found to be excitatory exactly corresponded to those classified as RS, while all inhibitory neurons were also FS. We only retained neurons spiking all throughout the recording for our analyses, yielding 71 neurons of which 21 were I neurons.

Similarly, spiking activity in layer III/IV of the pre-motor cortex of a macaque monkey was recorded by utah array, throughout a night, and an hour of target pursuit task on the following day. A 10-kHz sampling frequency was employed for recording. Classification of brain states was performed by visual inspection of the Local Field Potential (LFP), over time periods of 5 s, by identifying as SWS periods presenting large-amplitude oscillations in the 1-2 Hz frequency range. Spike-sorting yielded 152 neurons, of which 141 spiked throughout the whole recording. Clustering on features of the spike waveform has allowed for the sorting of neurons as putative E and I. Excluding neurons for which clustering was uncertain within a 30-percent margin yielded 81 neurons, of which 38 were I, over which all subsequent analyses were performed as presented in [11].

Time bin size was chosen in order to have one to few spikes from each neuron per time bin, while still having sufficient spikes per time bin from the whole population to compute statistics such as the pairwise covariances and the neuron-to-population dependencies. Since I neurons were consistently more active, this was equivalent to balancing a sufficient number of spikes from E neurons with sufficiently few spikes from any I neuron per time bin. In the human temporal cortex, where activity was considerably sparse, the chosen time bin size was 50 ms. In the interest of having comparable numbers of spikes per time bin and pairwise covariances, a time bin size of 25 ms was chosen for the monkey motor cortex, where firing rates were consistently higher than in the human temporal cortex.

## Appendix B: INFERENCE METHODS

Inferring the parameters from a MaxEnt model may be understood as a Lagrange multiplier problem, where one maximises the entropy, while taking as constraints that the desired model-predicted statistics match their empirical values. Then each model parameter is the Lagrange multiplier for one constraint, on one observable to reproduce. Taking, for example, the pairwise Ising model, the statistics we want to reproduce are the neuron mean firing rates and the pairwise covariances. The corresponding model parameters are the firing thresholds b_i_ and the pairwise couplings *J*_*ij*_ respectively. We therefore want to maximise

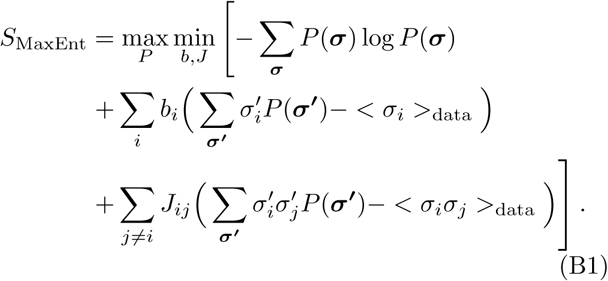

One can verify that maximizing *S*_MaxEnt_ with respect to *P* and with the chosen constraints [9, 15], gives the form of each of the models given previously in Eq. 1, Eq. 2, and Eq. 3. A Hessian analysis may prove that the problem is well-posed, as the solution exists and it is unique.

The key challenge thus resides in finding a method that quickly converges to this solution. Thanks to the explicit form of Eq. (B1), the gradient of the log-likelihood 
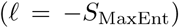
 with respect to the model parameters can be computed as differences between empirical and model-predicted averages of the conjugated observables. For example, the gradient with respect to the bias *b*_*i*_ of the Ising model can be estimated as 
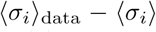
_model_. The inference can thus be performed by an ascendant dynamics that requires to estimate model averages of observables. For the Ising model we applied the Markov-Chain Monte-Carlo method introduced in [16, 26]. For the one population model, we applied the Newton dynamics proposed in [9] For the two-population model, we modify the algorithm of [9] to take into account two populations. We found that a simpler steepest descent dynamics, that does not take into account the Hessian, was fast enough for our data-sets.

## Appendix C: TUNING CURVE AND SENSITIVITY TO POPULATION

In order to quantify the dependence of each neuron on the rest of the population’s activity, we used tuning curves [9] and sensitivity to the population.

**Tuning curves** characterize the dependence of the average activity of a neuron conditioned to the activity of either the population, either the E or I sub-population. The tuning curve of neuron *i* is defined as 
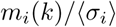
, where 
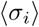
 gives the neuron’s mean activity across all time bins. *m*_*i*_(*k*), instead, denotes the neuron’s mean activity at fixed population activity and it is defined as:

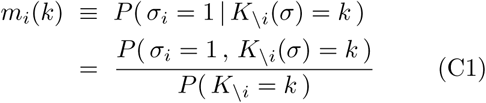

where 
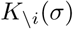
 is the number of active neurons in the configuration σ, when neuron *i* has been excluded.

**Sensitivity to population**. A tuning curve shows the whole profile of the dependence of a neuron activity on the rest of the population. In order to quantify this effect, we introduced the neuron sensitivity to the population, depicting the neuron’s fluctuation in activity across states of population activity: 

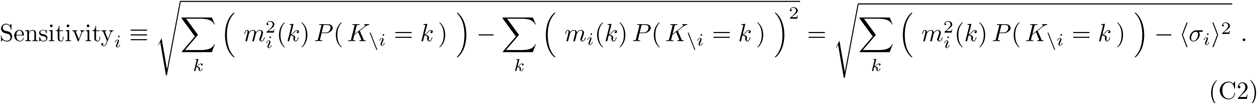

## Appendix D: TWO-POPULATION MODEL

In this section, we generalize the analysis of the one-population model introduced in [9] to the case of two populations. From our model introduced in Eq. 3, we can de ne the couplings 
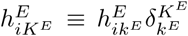
 for E neurons to the E and the I populations, and respectively for I neurons, such that the probability of a firing pattern occurring is 

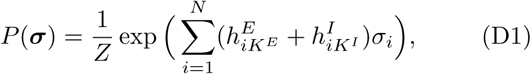

The model is said solvable as the normalisation *Z* can be expressed analytically. Note that the model is invariant under several gauge transformations as a number of linear combinations of its parameters 
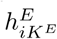
 and 
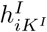
 do not a ect the probability distribution. Z and its derivative allow us to determine the statistics of the model, such as the mean firing rate and pairwise covariances.

**Normalisation**. From Eq. (D1) the normalisation is defined as 

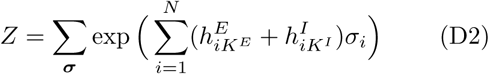

where we sum over all possible firing patterns σ. We may decompose this sum into terms 
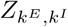
 with given E and I population activities, such that 

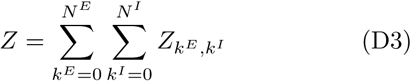

Then, we have 

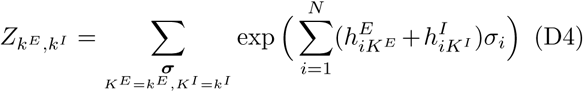

where we sum over all possible firing patterns for all neurons for which *K*^*E*^ excitatory neurons active and *K*^*I*^ inhibitory neurons active. This is equivalent to summing over all possible patterns of E and I neurons independently, i.e.: 

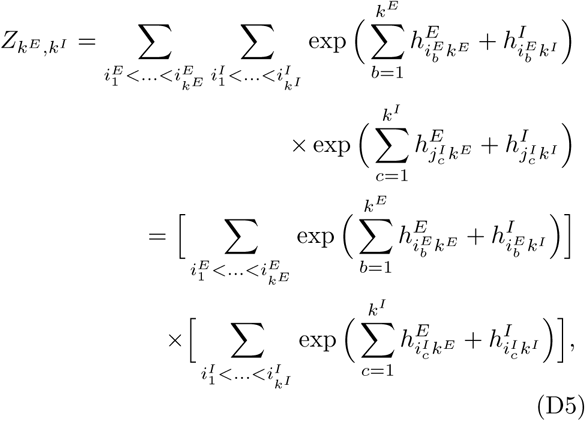

where the *i*^E^_b_ spans over all the active E neurons for a given E activation pattern, and respectively for the *j*_c_^I^ for active I neurons. The result may be written as a product of two terms as these terms share no parameters in common.

Here, the first term is summed over all possible firing patterns for E neurons that yield *K*^*E*^ = *k*^*E*^, and similarly for I neurons in the second term. Now, analogously to [9] let *Q* be a polynomial such that the products over all i 

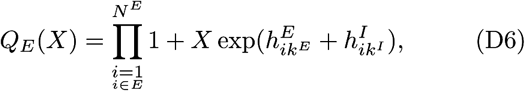

where we take the product over all *i* excitatory neurons, and similarly for *Q*_I_(*X*) multiplying over all inhibitory neurons. Now, the coefficient of *Q*_E_ of order X^k^
^E^, denoted 
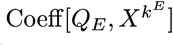
, corresponds to the sum over all the products of *k*^E^ terms of the form exp 
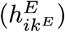
 or in other words, the sum over all products of combinations of E neurons 
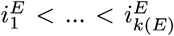
, which is exactly equivalent to the first term of equation D5. Since the same obviously applies for I neurons, we have

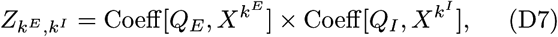

As the *Q* coefficients can be recursively computed, *Z* is analytically computable, thus the model is solvable. Next we derive the statistics of the model from *Z*.

**Model statistics**. The statistics predicted by the model is given by differentiating *Z*. We use this to predict the mean firing rates and pairwise covariances from the population couplings 
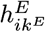
 and 
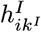
. As we defined in Eq.s (D2) and (D3), 

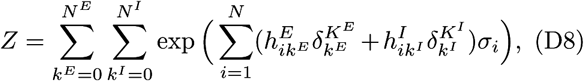

Thus, the joint probability of a given neuron spiking and a given number of E neurons spiking in any time bin is as follows: 

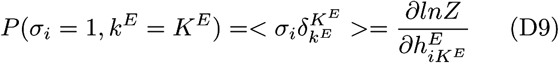

Recalling our expression for *Z*_*k*_^*E*^,_*k*_^*I*^ in Eq. (D7), this yields 

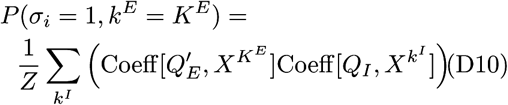

where 

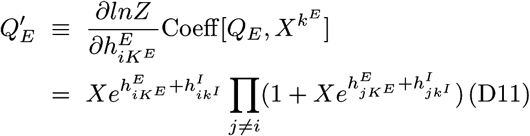

and *i* is an E neuron. For an I neuron’s firing probability one can swap around E and I in Eq. (D10). This allows the straightforward derivation of the firing rate by summing over all values of K^E^ (resp. *K*^I^ for I neurons).

Likewise, the pairwise correlations may be computed from 

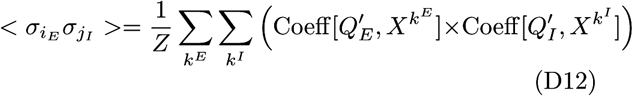

for two neurons of di erent types, and 

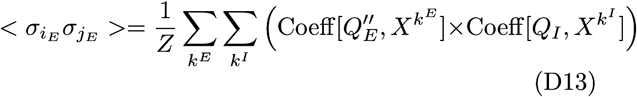

where 

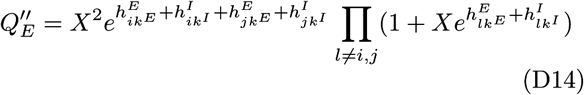

and i is an E neuron (extending to I neurons is very straightforward).

**Shuffle tests**. To verify whether information on neuron types signi cantly improves the model’s prediction performance, for each species, we perform a series of ten inferences on the same SWS data-set. Each time, the neuron labels are independently shu ed, while the number of E neurons and the number of I neurons remains the same. The Mean Square Error (MSE) on the predicted pairwise covariances is computed every time. We found it to be consistently larger for the shuffled trials compared to that where empirical neuron types are known. This is quantified by the Mann-Whitney *U* test on the samples of MSE_shuffled_[n] - MSE_data_ with n ranging from one to ten.

## Supplementarygures

**FIG. S1:**
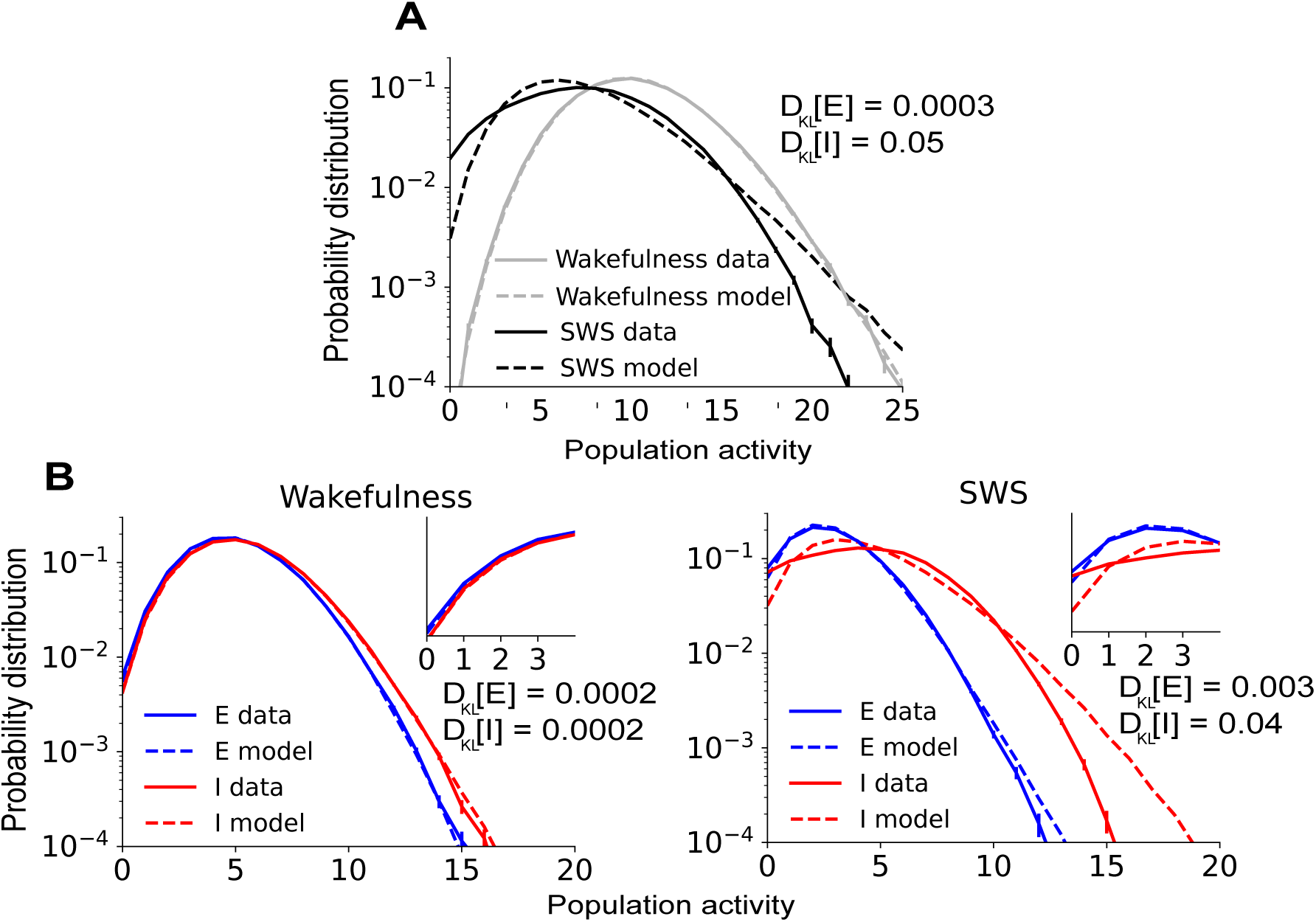
Pairwise Ising model analysis on monkey recording. **A**) Empirical and predicted distributions of the population activity *K* for the population of excitatory (E) neurons, and that of inhibitory (I) neurons. On the monkey data, the Ising also performs better at capturing the population statistics during wakefulness than SWS. **B**) Empirical and predicted population activities for E and I neurons. The model fails at reproducing the statistics of inhibitory population activity during SWS, similarly to with the human data.

**FIG. S2:**
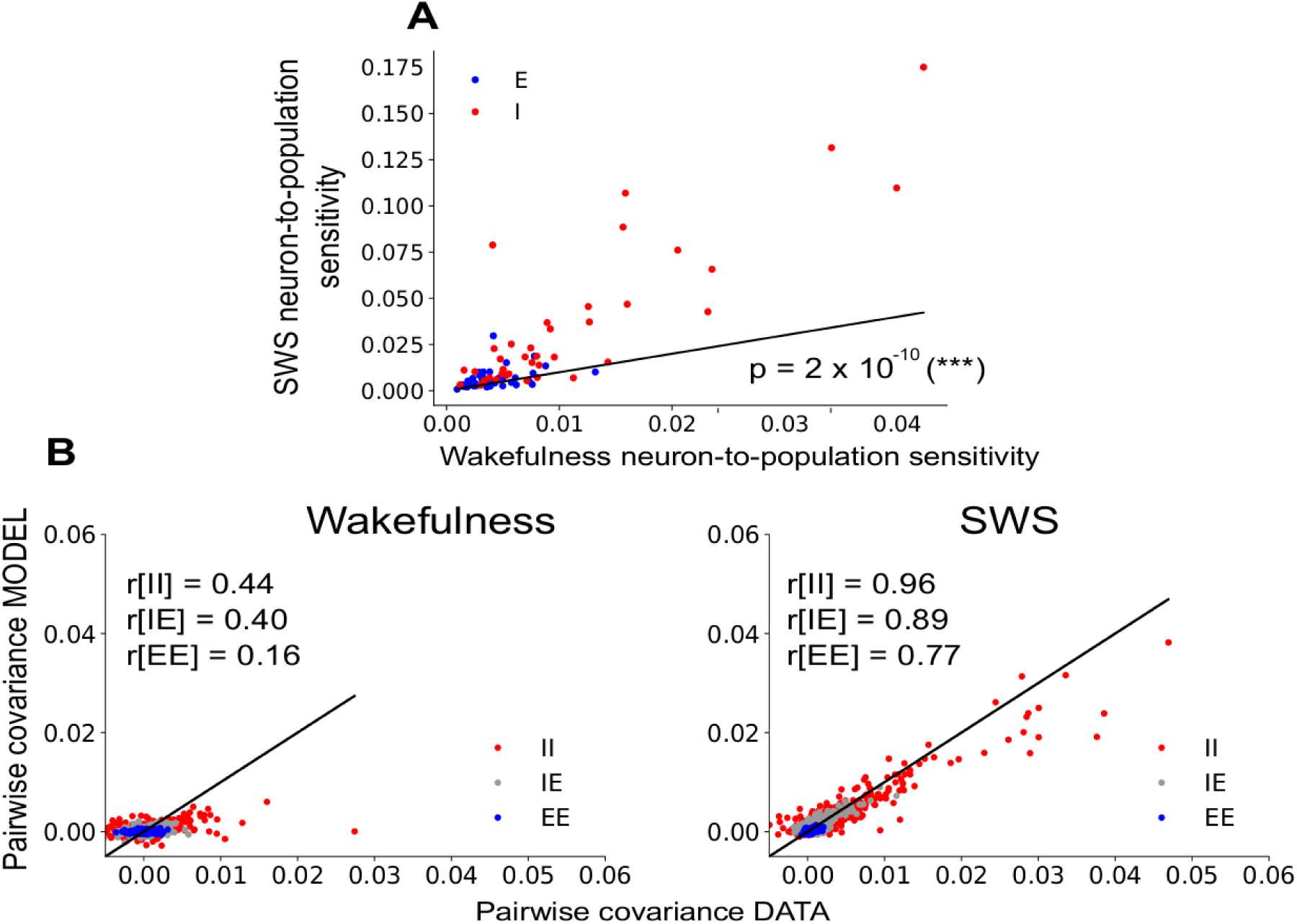
Single-population model analysis on monkey recording. **A**) Scatter-plot of the neuron sensitivity to the population activity in wakefulness and SWS. Neurons are consistently more sensitive during SWS (*p*-value <lt; 0:001). **B**) Pairwise covariances, empirical against predicted, for wakefulness (left) and SWS (right) states. Relative success for SWS, especially I-I pairs suggests these neurons are most responsive to whole-population activity, even though the model tends to under-estimate the larger pairwise covariances. The model completely fails to account for pairwise covariances during wakefulness.

**FIG. S3:**
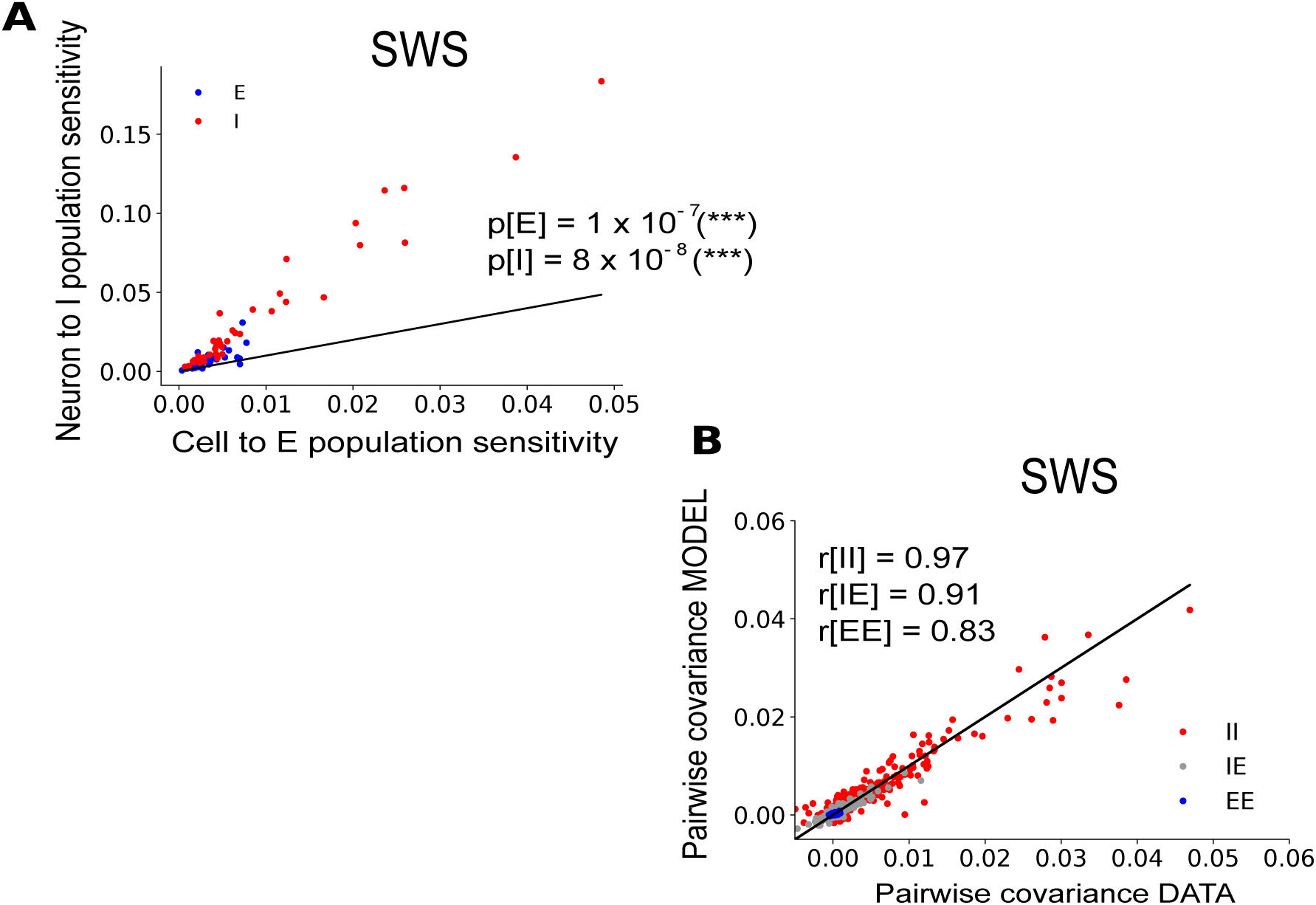
Two-population model shows significant improvement in prediction for all types of neurons. **A**) Scatter-plot of neuron sensitivity to E versus I population, during SWS. Both I and E neurons are more tuned to I population than the E one (*p*-value <lt; 10 ^-3^, Wilcoxon sign-ranked test). **B**) Pairwise covariances, empirical against predicted, for the two-population model, during SWS. Improvement compared to the whole-population model is confirmed by the Pearson correlations.

